# Deficits in decision-making induced by parietal cortex inactivation are compensated at two time scales

**DOI:** 10.1101/2021.09.10.459856

**Authors:** Danique Jeurissen, S Shushruth, Yasmine El-Shamayleh, Gregory D Horwitz, Michael N Shadlen

## Abstract

Perceptual decisions arise through the transformation of samples of evidence into a commitment to a proposition or plan of action. Such transformation is thought to involve cortical circuits capable of computation over time scales associated with working memory, attention, and planning. Neurons in the lateral intraparietal area (LIP) are thought to play a role in all of these functions, and much of what is known about the neurobiology of decision making has been influenced by studies of LIP and its network of cortical and subcortical connections. However a causal role of neurons in LIP remains controversial. We used pharmacological and chemogenetic methods to inactivate LIP in one hemisphere of four rhesus monkeys. Inactivation produced clear biases in decisions, but the effects dissipated despite the persistence of neural inactivation, implying compensation by other unaffected areas. Compensation occurs on a rapid times scale, within an experimental session, and more gradually, across sessions. The findings resolve disparate studies and inform interpretation of focal perturbations of brain function.

## Introduction

A decision is a commitment to a proposition or plan of action based on evidence from the environment or memory. The underlying neural computations convert such evidence into a state similar to working memory or motor planning. This conversion involves a network of brain areas spanning the association areas of the cerebral cortex as well as their subcortical connections. Even a simple decision to look to the left or right, based on visual evidence from straight ahead, is known to involve neurons in the dorsolateral prefrontal cortex, frontal eye field, striatum, superior colliculus, and lateral intraparietal area (LIP) (Shadlen and Newsome, 1996; Kim and Shadlen, 1999; Horwitz et al., 2004; Ding and Gold, 2010, 2012). Neurons in these areas represent both the saccadic choice and the evolving deliberative process—the integration of noisy evidence leading to the choice (Shadlen and Kiani, 2013).

The evidence-accumulation process has been extensively characterized in area LIP. Neurons in LIP combine accumulating evidence with other factors, including biases (e.g., prior probability) and time-costs to establish a representation (the decision variable) suitable for terminating the process. However, whether LIP, or any other single area, is essential to this process remains unclear. Causal perturbations of LIP activity have led to mixed results. Hanks et al. (2006) showed that electrical microstimulation of neurons that represent one of two choice targets caused a small bias in favor of that choice. The bias was associated with changes in response time by an amount consistent with a change in the firing rates of neurons that represent the decision variable. However, inactivation of LIP has not produced consistent effects on choice. Chen et al. (2020) observed striking biases against choice targets in the visual hemifield contralateral to cryo-inactivated posterior parietal cortex, including area LIP. However two recent studies used intraparenchymal infusions of the GABA-A agonist, muscimol, to inactivate LIP specifically, and found only small biases (Zhou and Freedman, 2019) or no behavioral effects at all (Katz et al., 2016).

We hypothesized that the weak behavioral effects might be explained by compensation from unaffected parts of the decision-making network (Fetsch et al., 2018). Such compensation could arise from neurons in distal brain regions (including the homologous LIP in the opposite hemisphere) as well as from local neurons within the targeted LIP but outside the inactivated region. We therefore inactivated LIP, but in contrast with previous studies, we (*i*) ensured that our inactivation encompassed a substantial fraction of the neurons that were associated with decision formation, and (*ii*) tracked the effect of inactivation over the course of each experimental session. We found that inactivation of area LIP induced a large bias in two types of perceptual decisions but only temporarily; the bias diminished within a few hundred trials and across inactivation sessions. The behavioral compensation was evident in monkeys performing two types of decision-making tasks, highlighting the generality of the phenomenon.

## Results

We trained four rhesus monkeys on perceptual tasks requiring a binary decision about a stimulus category. Monkeys 1 and 2 decided whether the net direction of random dot motion (RDM) was to the left or right (Fig. 1A). We varied the difficulty of the decision by controlling the strength and duration of the motion. After the removal of the motion stimulus, the monkeys reported the perceived net direction of motion with an eye movement to a choice-target on the right or left side of the display. Monkeys 3 and 4 made a decision about the temporal order of two flashed targets, which were presented sequentially in the left and right hemifield (Fig. 1B). Difficulty was controlled by the time between the onset of the two targets (Δ*t*). After a wait period following target presentation, the monkey had to report which of the two targets had appeared first by making an eye movement to the remembered location of that target.

**Figure 1:**
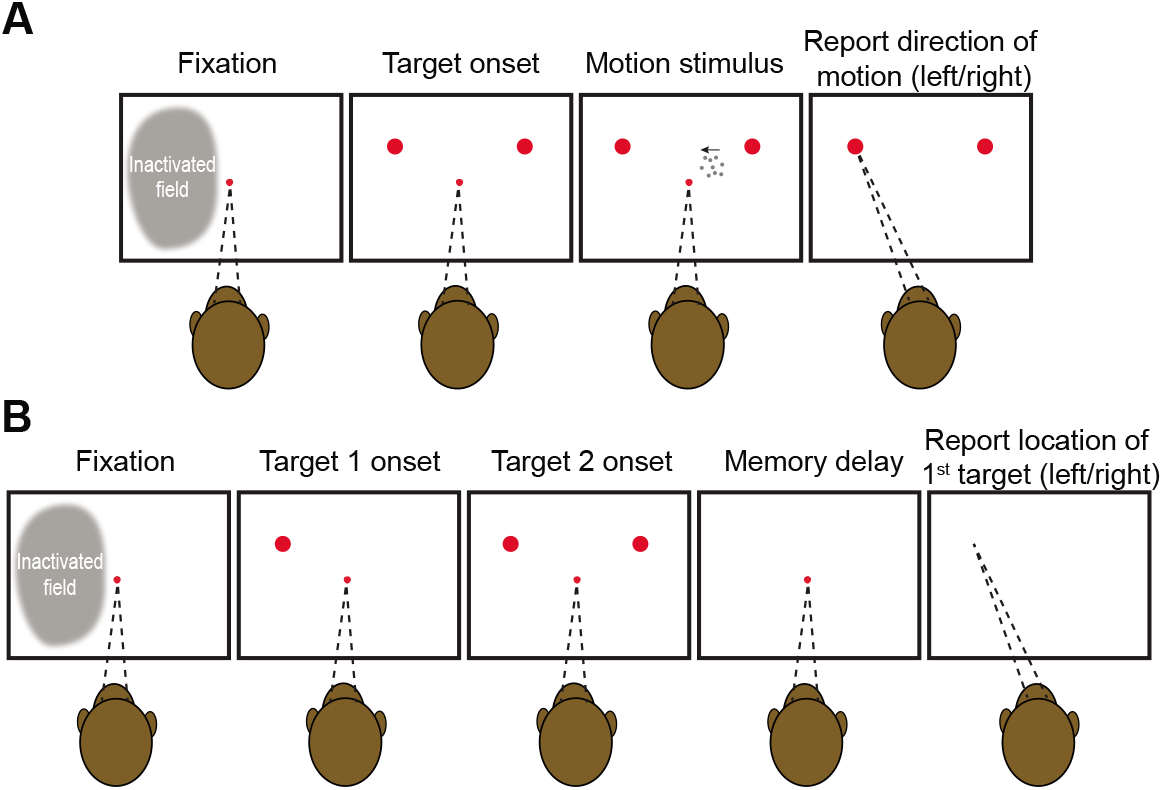
Behavioral tasks. Both tasks require the monkey to make a binary decision and report it with an eye movement to one of two choice-targets presented in the left or right hemifield, respectively. In each trial, the monkey is required to maintain its gaze on a central fixation point until its extinction, which serves as a *go cue*. **A**, Motion direction task. Dynamic random dot motion (RDM) appears within an invisible aperture contained within the hemifield ipsilateral to the inactivated cortex. The fixation point and motion stimulus are extinguished simultaneously, whereupon the monkey reports its decision. The monkey is rewarded for choosing the target in the direction of the motion (and randomly for the 0% coherent motion). Across trials, the strength, direction (left or right), and duration of the motion were varied randomly, as were the exact positions of choice targets (see Suppl. Fig. 2). **B**, Temporal order task. The choice-targets are presented sequentially. Choice-targets 1 and 2 are extinguished simultaneously, 430 ms after the onset of the first target. The fixation point is then extinguished after a variable delay, and the monkey is rewarded for making a saccade to the remembered location of the first target. Across trials, the order, onset asynchrony, and exact positions of the targets were varied randomly.

The two tasks share the requirement of reporting the decision with an eye movement. In such tasks, neurons in area LIP that exhibit spatially selective persistent activity during saccade planning (Gnadt and Andersen, 1988) are thought to play a role in decision formation (Shadlen and Newsome, 1996; Wardak et al., 2002; Rorie et al., 2010). We used a memory-guided saccade task (Gnadt and Andersen, 1988) to ascertain the full extent of LIP (in one hemisphere) that contains such neurons. Consistent with previous reports (Patel et al., 2010), neurons with persistent activity were identified across a broad swath of the lateral bank of the intraparietal sulcus (IPS). The anteroposterior spread ranged from 6–10 mm; the dorsoventral spread ranged from 3–7 mm (Fig. 2A–B). We targeted our inactivation to the region determined by this functional mapping in each monkey. In Monkeys 1–3, we inactivated the region of interest by making several injections of the GABA-A agonist muscimol. In Monkey 4, we injected an AAV vector to express the inhibitory muscarinic receptor hM4Di in the region of interest (Armbruster et al., 2007) and targeted the receptor by subcutaneous administration of clozapine. We confirmed that our inactivation encompassed the targeted area by multi-neuron recordings (Fig. 2C-D).

**Figure 2:**
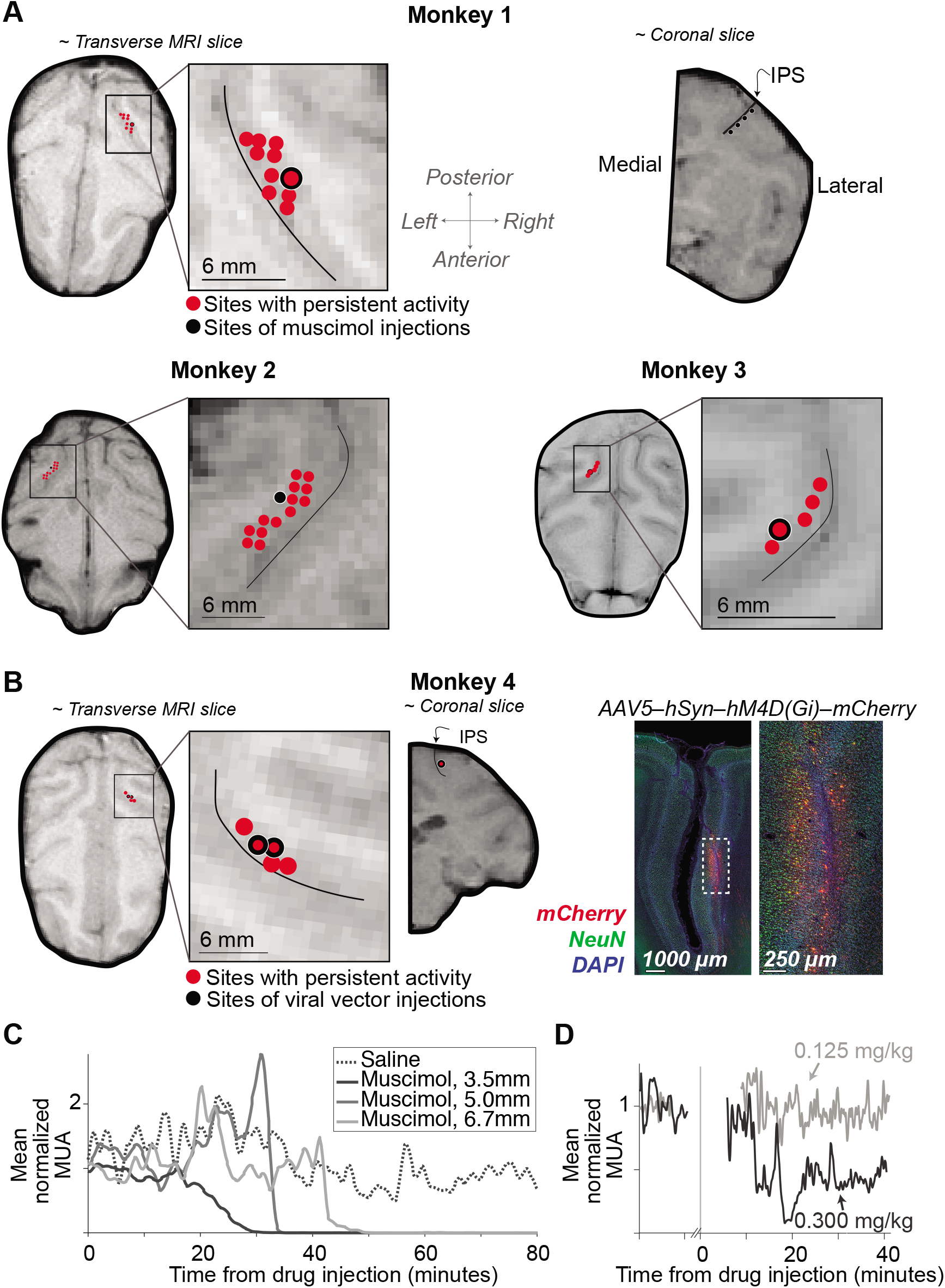
Localization and characterization of LIP inactivation sites. **A**, The locations of muscimol injections and neurons with spatially selective persistent activity are superimposed on MRIs for Monkeys 1, 2, and 3. The near-transverse planes are orthogonal to the injection trajectories. The nearcoronal MRI slice from Monkey 1 (top-right) shows positions along the intraparietal sulcus (IPS) where muscimol was injected. The thin black curve (*inset*) marks the center of the IPS. **B**, *Left*: Location of viral vector injections for Monkey 4. The red points in the MRI are sites containing neurons with spatially selective persistent activity. The coronal slice shows the injection site; same conventions as in *A. Right*: Representative histology. Expression of hM4Di-mCherry receptor is restricted to the lateral bank of the IPS. **C**, Time course of multi-unit activity (MUA) in area LIP following injection of saline (dotted) and muscimol (solid). Recordings were obtained at different distances from the injection site (legend). Note the complete suppression of activity in < 1 hour. **D**, Time course of MUA following subcutaneous injection of clozapine at the lowest (gray) and highest (black) dose tested.

In both tasks, the choice targets were in opposite hemifields, contralateral and ipsilateral to the inactivated area LIP. We refer to the corresponding choices as contraversive and ipsiversive, respectively. By convention, positive values of motion strength (task 1) and target asynchrony (task 2) indicate evidence for the contralateral choice target. Figure 3 shows the choice behavior over the first 100 trials of the first LIP inactivation experiment for each monkey. The rationale for restricting analysis to the earliest trials and sessions will be made clear in Figure 4. All monkeys made fewer contraversive choices during the 100 trials after inactivation than they did during the 100 trials before inactivation. This reduction held at nearly every stimulus strength in all four monkeys (Fig. 3). Thus the monkeys made more errors in response to contraversive motion (Fig. 3A) and early contralateral target appearances (Fig. 3B). The effect could not be attributed to more frequent fixation breaks on trials supporting a contralateral choice compared to an ipsilateral choice (Fisher exact test, *p* > 0.15 for each monkey). The sigmoid curves in Figure 3 are fits of a logistic regression model (Eq. 5). The fits show clear effects of muscimol and hM4Di mediated inactivation on the monkeys’ decisions compared to pre-inactivation and to control experiments. The dominant effect of inactivation is a bias against contraversive choices (*p* < 0.02 in all cases, Table 1). Inactivation also appears to affect the slope of the choice functions, which would suggest decreased sensitivity to motion (Monkey 1) and Δ*t* (Monkeys 3 & 4). The effect is statistically significant in Monkey 3 (*p* < 0.01, Table 2), and it is statistically significant in Monkey 1 upon inclusion of more experimental sessions (Eq. 9, *p* < 0.023). Overall, however, we interpret the the effect of inactivation on sensitivity as inconsistent across animals, and therefore inconclusive. From here on we focus all analyses on the decision bias.

**Figure 3:**
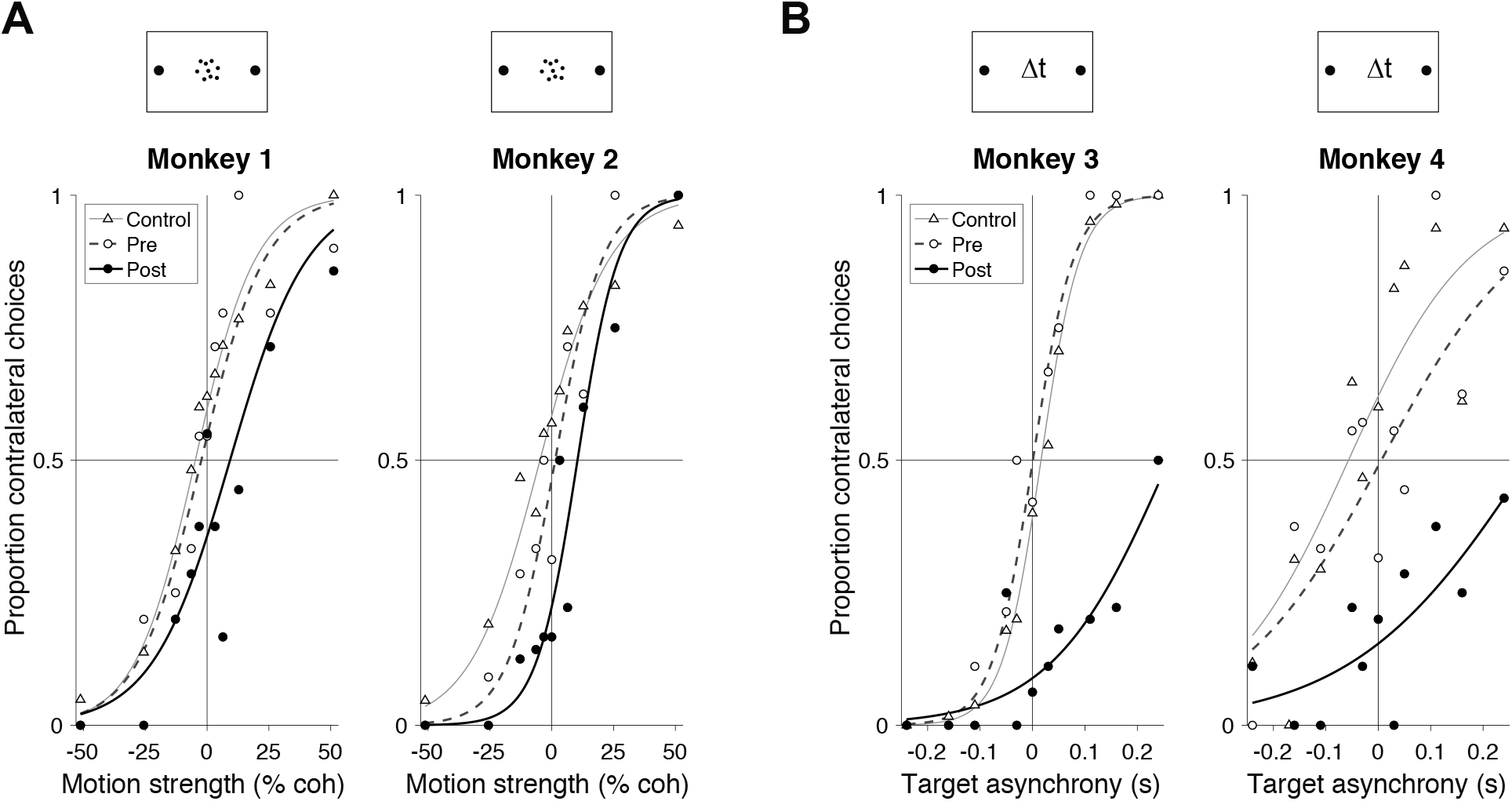
Inactivation of LIP induces a decision bias. **A**, Proportion of contralateral choices as a function of motion strength for Monkeys 1 and 2. Filled circles show data from the first 100 trials after muscimol injection in the first inactivation session. Open symbols show data from the last 100 trials in the pre-injection phase of the same experiments. Triangles depict data from all control sessions using the first 100 trials after the saline or sham injection. Muscimol induces a bias against contralateral choices. Curves are logistic regression fits (Eq. 2). **B**, Proportion of contralateral choices as a function of target onset asynchrony for Monkeys 3 and 4. Data from Monkey 3 are from the first session in which muscimol was administered. Data from Monkey 4 are from the session in which 0.3 mg/kg clozapine was administered. Other conventions are the same as in *A*.

**Figure 4:**
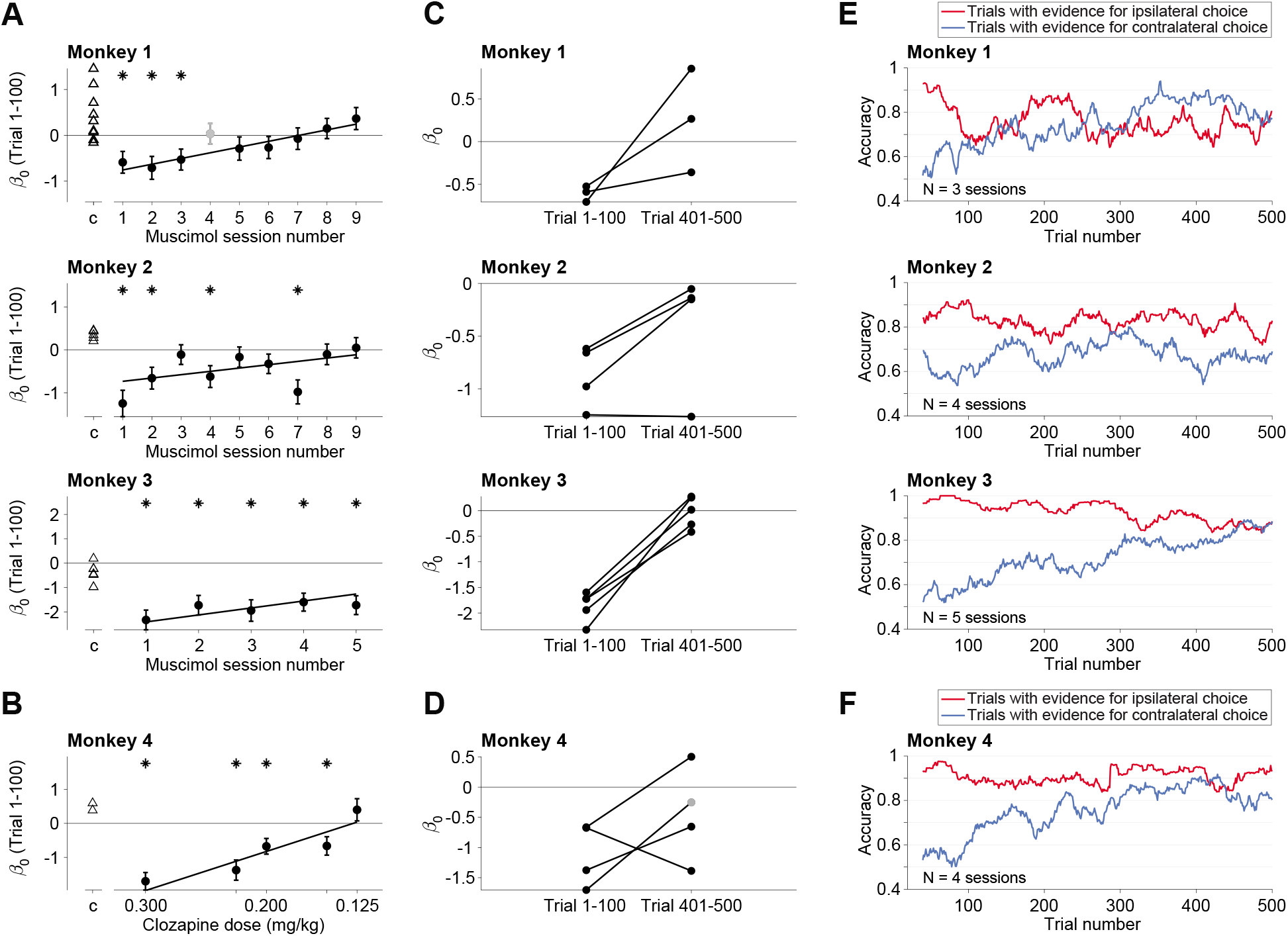
Compensation of bias across and within sessions. **A**, The size of the contraversive bias (*β*_0_, Eq. 2) in the first 100 trials following inactivation is plotted as a function of experimental session. Data are shown separately for the three monkeys that received muscimol. Negative bias (*β*_0_ < 0) indicates bias against contraversive decisions. Triangles are data from control sessions. Asterisks denote statistical significance (*p* < 0.05; 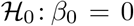). The regression line is from the fit to Eq. 6, excluding session 4 for Monkey 1 (gray point), in which we appraised a smaller volume (8 *μL*; see Methods). Error bars are s.e. **B**, Effect of clozapine dose on decision bias (Monkey 4). Same conventions as in *A*. **C–F**, Within session compensation. These analyses use only sessions with a statistically significant bias in the first 100 trials (asterisks in A & B). **C**, Individual muscimol sessions. Each line connects the bias in trials 1–100 with the bias in trials 401–500. **D**, Individual clozapine sessions (Monkey 4). Same conventions as in C, except for one session, where less than 500 trials were completed. The gray point is the bias from the last 100 trials (trials 186–286). **E,F** Gradual diminution of the bias. These analyses combine the individual experiments in C & D and group trials with the same sign of evidence (color), regardless of evidence strength (trials with 0% coh or Δ*t* = 0 are excluded). The traces are running means of choice accuracy using 40 trials. Trial numbers on the abscissa correspond to the end of the averaging window.

**Table 1:** *β*_2_ values, SE and *p* values from Eq. 5

|  | pre vs. post |  |  | control vs. post |  |  |
| --- | --- | --- | --- | --- | --- | --- |
| | value | SE | $p$ | value | SE | $p$ |
| Monkey 1 | -0.77 | 0.34 | 0.02 | -0.98 | 0.25 | 9e-05 |
| Monkey 2 | -1.08 | 0.40 | 0.01 | -1.58 | 0.32 | 1e-06 |
| Monkey 3 | -2.31 | 0.50 | 4e-06 | -1.90 | 0.42 | 6e-06 |
| Monkey 4 | -1.66 | 0.37 | 6e-06 | -2.20 | 0.34 | 1e-10 |

**Table 2:** *β*_3_ values, SE and *p* values from Eq. 5

|  | pre vs. post |  |  | control vs. post |  |  |
| --- | --- | --- | --- | --- | --- | --- |
| | value | SE | $p$ | value | SE | $p$ |
| Monkey 1 | -0.02 | 0.03 | 0.58 | -0.03 | 0.03 | 0.32 |
| Monkey 2 | 0.03 | 0.06 | 0.63 | 0.07 | 0.05 | 0.13 |
| Monkey 3 | -16.68 | 6.61 | 0.01 | -17.77 | 3.87 | 4e-06 |
| Monkey 4 | -1.38 | 3.11 | 0.66 | -2.79 | 2.78 | 0.32 |

Inactivation with muscimol reduced contraversive choices in the first session, but this effect diminished over subsequent sessions. Figure 4A shows the bias during the first 100 trials in each muscimol session compared to controls. All three monkeys exhibited weaker biases against contraversive choices in later sessions. For monkeys 1 (motion) and 3 (time), the change is strikingly monotonic (*p* = 10^-4^ and 0.003, respectively; Eq. 6, 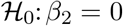). The trend is not monotonic in Monkey 2, but the decrease as a function of session number is statistically significant (*p* = 0.01). This effect is not explained by decreased efficacy of muscimol across sessions, as the drug induced silencing of neural activity in all sessions. Thus the decision-making network can learn to compensate for the loss of area LIP across multiple days. For Monkey 4, we varied the dosage to the agonist, clozapine, across sessions. As shown in Figure 4B, the contralateral bias was strongly dose dependent (*p* = 10^-8^). We did not detect an effect of session number in Monkey 4 (*p* = 0.15), possibly because it was masked by a strong effect of drug dosage, which was randomized across sessions. We cannot ascertain whether the lack of across-session compensation is attributed to the chemogenetic approach, or the limited number of sessions possible in this monkey, or the confounding effect of clozapine dose.

In addition to the behavioral compensation observed across sessions, the bias also dissipated over the course of individual sessions. In most sessions, the bias decreased gradually over a few hundred trials and resolved nearly completely after 500 trials (Fig. 4C–D). Figure 4E–F highlights this within-session attenuation of bias by combining sessions in which a statistically significant bias was present in the first hundred trials (asterisks in Fig. 4A, B). The initial bias is evident in the small fraction of correct contraversive choices (~60%) and the large fraction of correct ipsiversive choices (80–100%). This assay for the bias ignores stimulus strength, but it allows us to focus on the effect of trial number within a session by combining over strengths and sessions. The running means thus reveal a gradual dissipation of the disparity between accuracy on the contralateral and ipsilateral supporting stimuli. These changes were highly reliable by logistic regression for three of the monkeys (*p* < 10^-5^; Eq. 8, 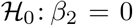) and borderline for Monkey 2 (*p* = 0.06).

The behavioral compensation across trials was not caused by recovery of neural activity at the inactivated site, which persisted for the entire duration of each session (Fig. 2C–D). Further, the monkeys still displayed signs of contralateral hemineglect on a simple extinction (side-preference) assay (Christopoulos et al., 2018a), conducted at the end of the experiment (Monkeys 2 and 3). Both animals exhibited a strong bias for choosing the treat presented in the ipsilateral visual field (compared to control sessions, *p* < 10^-3^, for both monkeys, Fisher exact test, Supp. Fig. 1). Thus the compensation exhibited on the perceptual decisions appears to be task specific.

## Discussion

We have shown that suppression of neural activity in cortical area LIP induces behavioral changes in perceptual decision making. We used two types of tasks and two methods of inactivation. In all cases, inactivation of LIP in one hemisphere produced a bias against contralateral choices, consistent with partial spatial hemineglect (Mattingley et al., 1998). The effect was transient, however, bringing to light compensatory mechanisms that operate on at least two distinct time scales—over the course of a few hundred trials within individual sessions and across multiple sessions separated by days. Our results complement a previous study that reported an even faster, within-trial compensation, associated with optogenetic suppression of neurons in extrastriate cortical area MT/V5 (Fetsch et al., 2018).

Previous studies have shown that unilateral inactivation of LIP produces behavioral effects consistent with contralateral hemineglect (Li et al., 1999; Chafee and Goldman-Rakic, 2000; Li and Andersen, 2001; Christopoulos et al., 2018b), and it affects target selection in attentionally demanding tasks (Wardak et al., 2002, 2004). Our findings complement these studies by showing that LIP inactivation affects decisions in a manner similar to a change in base-rate, prior probability, or value difference (Hanks et al., 2011; Rorie et al., 2010; Platt and Glimcher, 1999). The findings are also consistent with the observation that electrical microstimulation of LIP biases decisions regarding random dot motion in favor of contraversive choices (Hanks et al., 2006). Indeed the present findings would be unsurprising were it not for (*i*) the accompanying compensation and (*ii*) two recent studies that reported such a bias to be absent (Katz et al., 2016) or vanishingly small (Zhou and Freedman, 2019). The present findings readily explain this discrepancy.

We attempted to inactivate the extent of area LIP that contains neurons with spatially selective persistent activity during the saccade planning phase of an oculomotor delayed response—neurons that have been shown to represent the accumulation of evidence during perceptual decision making. Our mapping protocol revealed that the span of such neurons is extensive, consistent with Patel et al. (2010), indicating that a large injection of muscimol would be required to inactivate most of them. Thus the volume of cortex inactivated in our experiments was approximately 1.5 times the volume inactivated by Zhou and Freedman (2019) and Katz et al. (2016). We suspect that only a fraction of the relevant neurons were silenced in those studies, leaving open the possibility of weaker effects and more rapid compensation by neurons in the penumbra of the silenced tissue. Additional differences between these previous studies and ours may also contribute to the difference in results, including the levels of difficulty, jittering of the target positions, or differences in the motion display itself (e.g., highly salient moving elements in the Katz et al. study) which may discourage integration of evidence over time. Unless the animal is integrating information over time, relying on working memory (Constantinidis et al., 2018), or evaluating an interval of time itself (e.g., Leon and Shadlen 2003), there is little reason to expect LIP to play a role in the decision.

The finding of compensation has broad implications for the interpretation of causal studies. Otchy et al. (2015) showed that successful inactivation experiments (i.e., leading to loss of function) need not implicate the brain tissue targeted by the causal intervention—related to the concept of *diaschisis* in neurology (Carrera and Tononi, 2014). Our finding adds the complementary caveat that inactivation experiments yielding negative results do not rule out a causal role of the inactivated tissue. In other words, causation does not imply necessity. Yet, the phenomenon of compensation is likely to play a more constructive role in neuroscience than muddying the interpretation of null inactivation experiments. Translational neuroscience stands to benefit greatly from a fuller characterization of behavioral compensation and its underlying mechanisms. The present study and Fetsch et al. (2018) only begin to scratch the surface.

Rapid compensation would seem to rely on mechanisms of plasticity that operate on behaviorally relevant time scales (e.g., Magee and Grienberger 2020). One possibility is, shortly after LIP inactivation, downstream areas sense that the source of information they rely upon is compromised and establish communication with alternate sources. The mechanisms underlying such flexible routing of information from the senses to circuits that control behavior is unknown. Yet they are essential for higher brain function, for which dedicated input-output relations were not anticipated by evolution and therefore not determined by dedicated pathways. We suspect that these mechanisms involve both long range cortico-cortical feedback and matrix thalamic projections to superficial cortical layers (e.g., Jones 2001). The same mechanisms might underlie the resiliency of humans to focal cortical lesions (Cramer et al., 1997)—the clinical observation that small strokes are often silent until there are enough of them (e.g., vascular dementia). So the news is mixed: on the one hand, the possibility of compensation exposes the limitations of causal manipulation to assign cognitive functions to localized regions of the brain. On the other, causal manipulations might be used to investigate the mechanism of compensation and to augment them to achieve clinically relevant goals.

## Acknowledgements

We thank Anthony Napolitano, Brian Madeira, Cornel Duhaney, Lee Anthony Williams, and the Institute of Comparative Medicine for technical support and animal care; John Neumaier of University of Washington for advice on chemogenetic techniques; and members of the Shadlen lab for helpful discussions.

The research was supported by the Howard Hughes Medical Institute; an R01 from the NIH Brain Initiative (MNS, R01NS113113); an R21 from the NIH National Institute on Aging (MNS, DJ, SS, 1R21AG067108-01); a postdoctoral fellowship from the Simons Collaboration on the Global Brain (Simons Foundation, DJ, 414196); a pilot grant from the Alzheimer’s Disease Research Institute at the Taub Institute, Columbia University (MNS, DJ, SS); Young Investigator Awards from the Brain and Behavior Research Foundation (DJ, 28476 and SS, 23556), and an R01 grant from the National Eye Institute (GDH, EY030441).

## Materials and Methods

All training, surgery, and experimental procedures were conducted in accordance with the Public Health Service Policy on Humane Care and Use of Laboratory Animals (National Research Council, 2011). Experiments were approved by the Columbia University Institutional Animal Care and Use Committee (IACUC) under protocol number AC-AAAW4454.

### Subjects

We performed extracellular neural recordings and unilateral reversible inactivation in the parietal cortex of four adult male rhesus macaques. The animals weighed 10, 7, 10, and 8 kg, and were aged 9, 18, 18, and 12 years, respectively. We used a pharmacological approach for inactivation in Monkeys 1, 2, and 3 and a chemogenetic approach in Monkey 4. All four monkeys had a headpost to allow head fixation and a CILUX recording chamber (Crist Instruments) over the parietal cortex. Recording chambers provided access to the right hemisphere in Monkeys 1 and 4 and to the left hemisphere in Monkeys 2 and 3.

### Behavioral Tasks

Visual stimuli were presented on a CRT monitor (60 or 75 Hz refresh rate; viewing distance 58 or 48 cm). Eye position was recorded using an infrared eye tracker (Eyelink, SR Research; sampling rate: 1 kHz). Stimuli were generated using the Psychophysics Toolbox (Brainard, 1997) in Matlab (Mathworks) under the control of a REX system (Hays Jr et al., 1982). Juice rewards were delivered by a solenoid-based reward system.

### Motion direction task

Monkeys 1 and 2 were required to decide whether the net direction of motion in a dynamic random dot display was leftward or rightward (Fig. 1A). The animal initiated each trial by fixating within ±4 degrees visual angle (dva) of a central red fixation point on a black background. After 0.6–1 s, two red choice-targets appeared in the left and right upper quadrants of the visual field. The exact location of each target was chosen randomly and independently on each trial using a uniform distribution of polar angle and eccentricities within a specified range (See Supp. Fig. 2). We took this step to ensure that the monkey could not infer the location of one target from the position of the other. After a random wait duration (drawn from a truncated exponential distribution, range 0.8–1.5 s, mean 1 s), the RDM stimulus appeared within a circular aperture (radius: 2.5 dva), at an eccentricity of 3.5 dva from the fixation point. The RDM was confined to the hemifield ipsilateral to the inactivated LIP. The RDM was generated using previously described methods (Roitman and Shadlen, 2002). Three interleaved sets of dots (density 16.7 dots/deg^2^/s) were presented on successive video frames. Each dot was redrawn three video frames later at a random location within the stimulus aperture or at a location consistent with the direction of motion; the motion coherence is the probability of the latter occurring. The coherence on each trial was drawn randomly from the set ±[0, 0.032, 0.064, 0.128, 0.256, 0.512]. Positive values indicate that the motion was towards the target in the hemifield contralateral to the inactivation site; negative values indicate motion towards the target in the ipsilateral hemifield. On 0% motion coherence trials, one of the targets was randomly assigned as correct. The RDM was presented for a variable duration drawn from a truncated exponential distribution (range 0.1–2 s):

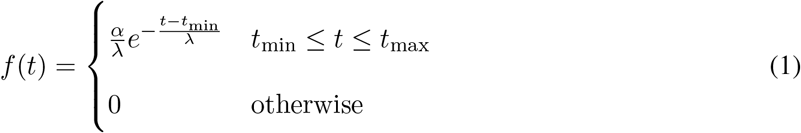

where λ = 0.3, *t*_min_ = 0.1 s, *t*_max_ = 2 s, and *α* is chosen to ensure the total probability is unity. The fixation point and RDM disappeared simultaneously, whereupon the monkey was allowed to indicate its decision about the direction of motion by making a saccade to the corresponding target.

For Monkey 1, we used a fixed ratio reward schedule with a juice reward for every correct trial. For Monkey 2 we used a variable ratio reward schedule with a juice reward for only a subset of the correct trials. The number of correct trials needed to obtain a reward was a random number drawn from a Normal distribution, 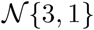, and discretized to the nearest integer from 1–6. Incorrect trials were never rewarded and were followed by a time-out (5 s).

### Temporal order task

Monkeys 3 and 4 performed a temporal-order discrimination task in which they indicated which of two targets appeared first (Fig. 1B). The animal initiated a trial by acquiring a central red fixation point. After 0.6–1 s of maintained fixation, two targets appeared, one in each hemifield at locations that were randomized across trials, as in the motion task (Supp. Fig. 2). The delay between targets were randomly chosen on each trial from the set ±[0, 27, 53, 107, 160, 240] ms, where positive values indicate that the target contralateral to the inactivated side was presented first. The first target stayed on the screen for 0.43 s, and both targets disappeared simultaneously. Following a memory delay (drawn from a truncated exponential distribution, range 1–2 s, mean 1.4 s), the monkey was required to make a saccade to the location of the remembered target that had appeared first to obtain a juice reward. Both monkeys were rewarded using a fixed ratio reward schedule with a juice reward for all correct trials and on 50% of the trials in which the targets appeared simultaneously.

### Side-preference test

Monkeys 2 and 3 were tested for signs of spatial hemineglect (Christopoulos et al., 2018a) at the end of experimental sessions. The testing was performed after retraction of any pipettes and electrodes in the brain but before the head fixation was released. Two equal sized pieces of fruit were offered to the monkey, one to the left and another to the right, equidistant from the mouth. The animal indicated its choice for one piece of fruit by extending its tongue to one side or the other to acquire that treat. This procedure was repeated 8–16 times per session. A Fisher exact test was used to compare the proportion of ipsilateral choices between tests conducted after inactivation sessions and after control sessions (Supp. Fig. 1). On interleaved control trials, a single piece of fruit was offered unilaterally to confirm that the monkey could indicate choices on both sides.

### History of participation in previous causal manipulation experiments

Three of the monkeys had participated in other causal manipulation experiments. We provide details here for completeness. Monkeys 1 and 2 had participated in an experiment in which small clusters of cells in area MT were inhibited using optogenetics (Fetsch et al., 2018) or stimulated using electrical stimulation (Fetsch et al., 2014) in a post-decision wagering task. Before training on the temporal-order discrimination task and before the injection of the viral vectors, Monkey 4 participated in 5 sessions in which we optimized our muscimol infusion techniques. During these sessions, muscimol was infused into area LIP while the monkey performed simple saccadic tasks.

### Pharmacological inactivation and neural recordings

We used magnetic resonance imaging (MRI) to localize the intraparietal sulcus (IPS) in relation to the recording chamber. We obtained MR images (T1 weighted gradient-echo sequences in Monkeys 1, 2, and 4; a T2 weighted spin-echo sequence in Monkey 3) with a recording grid *in situ*. We used custom software to project the recording grid onto the MR images (Fig. 2A,B). We systematically mapped the lateral bank of the IPS and noted the locations of neurons with spatially selective persistent activity during visually-guided and memory-guided saccade tasks (Gnadt and Andersen, 1988). We planned our inactivation to encompass as many of these locations as possible.

Muscimol and saline injections were made with quartz glass injection pipettes (115 μm outer diameter, 85 μm inner diameter, beveled tip, Thomas Recording). Extracellular neural recordings were obtained with a tungsten microelectrode (100 μm outer diameter, ≈1 *M*Ω impedance, FHC Inc.) to confirm tissue silencing Fig. 2C) and to estimate its spatial extent. The pipette and the microelectrode were advanced independently using a motorized hydraulic drive (Narishige International Inc.) along parallel trajectories through the IPS. The mean distance between the electrode and first injection site was 3.6 mm (range of 2.1—6.7 mm across sessions). A grid system allowed us to place the pipette at a site with an abundance of the targeted neurons and sufficiently near other targeted sites to achieve inactivation by diffusion from multiple injections spaced along this single trajectory (see Fig. 2A and Table 3). The injection site and depths were the same in all sessions for a given monkey.

**Table 3:**
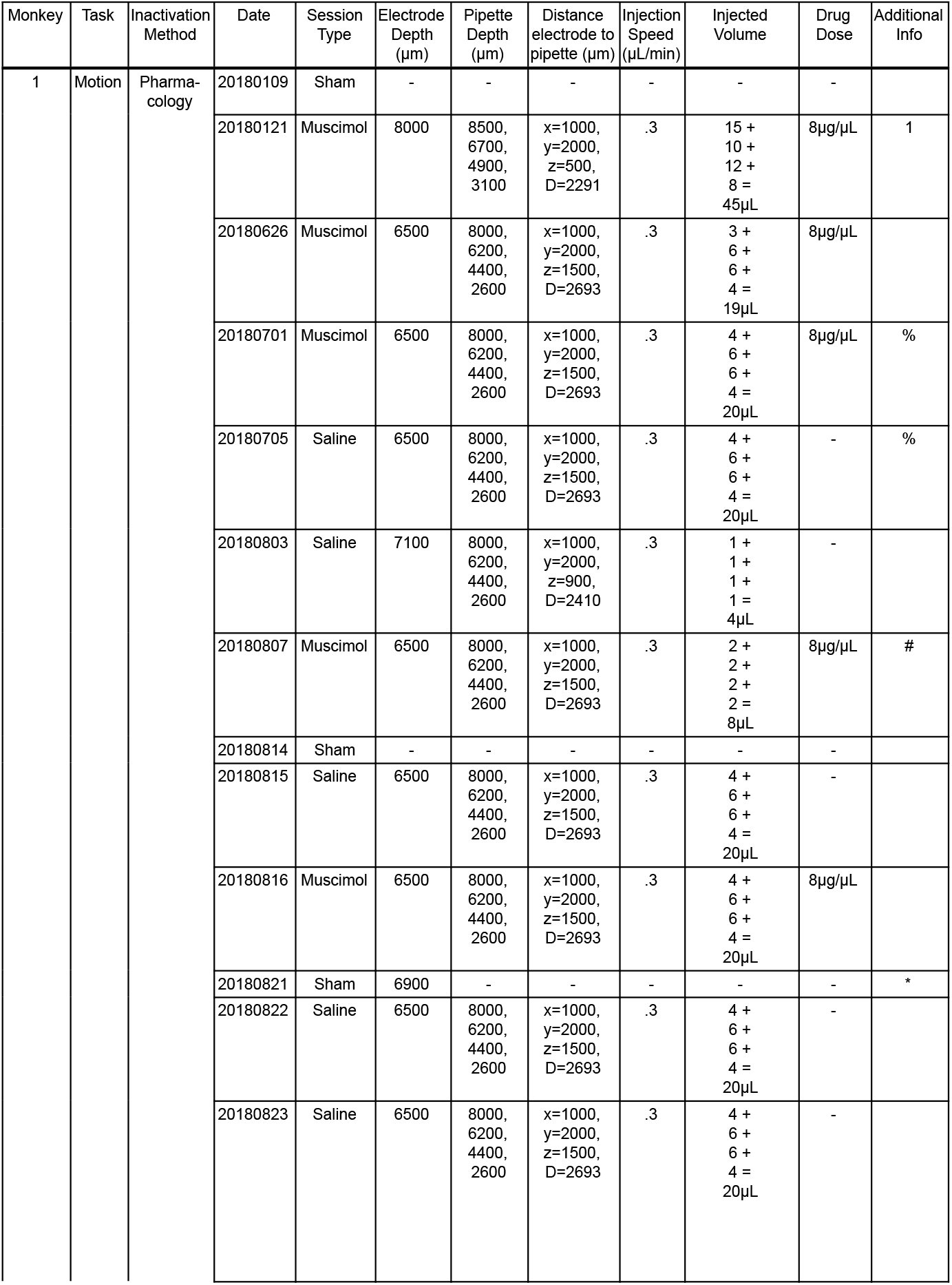
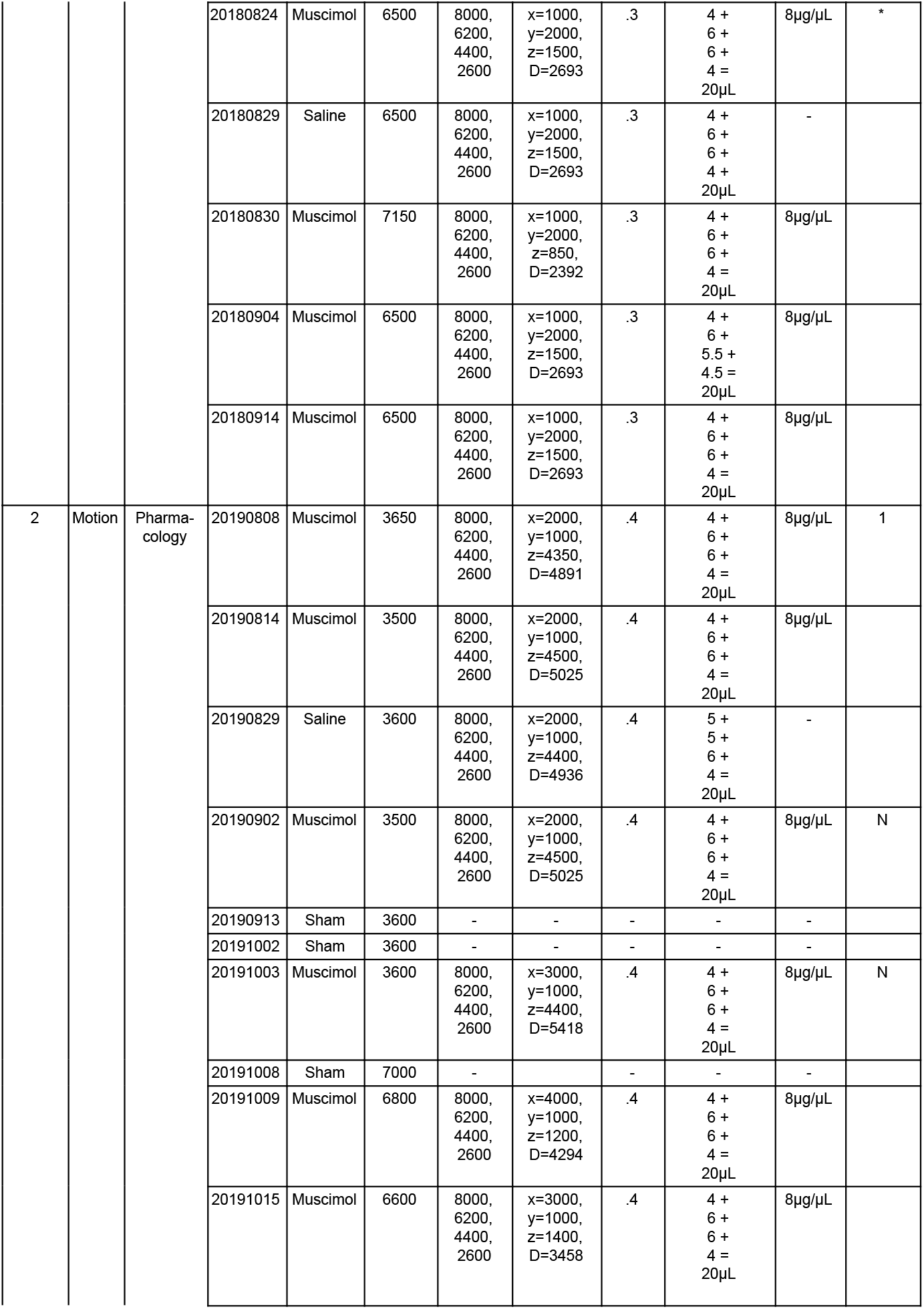
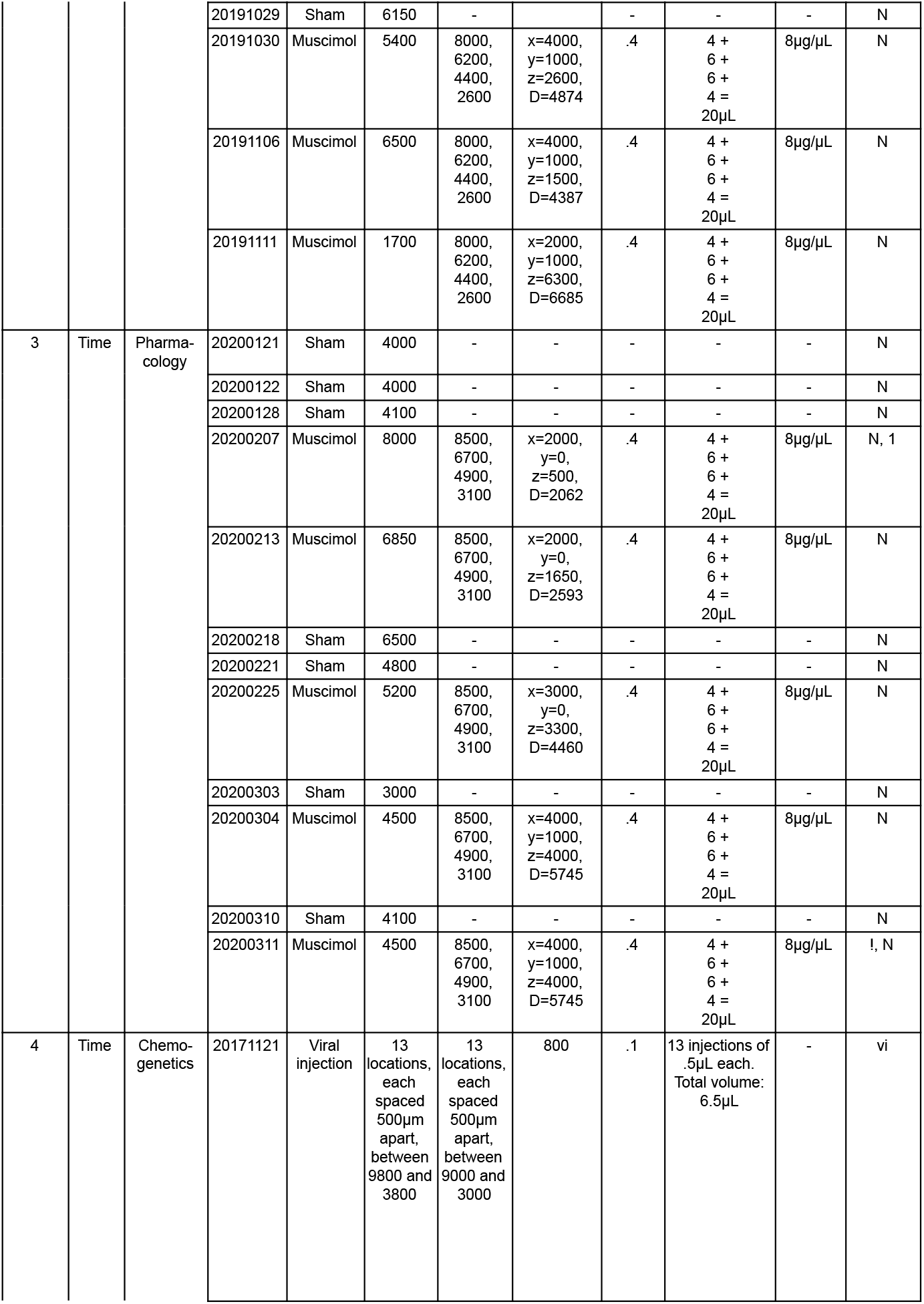
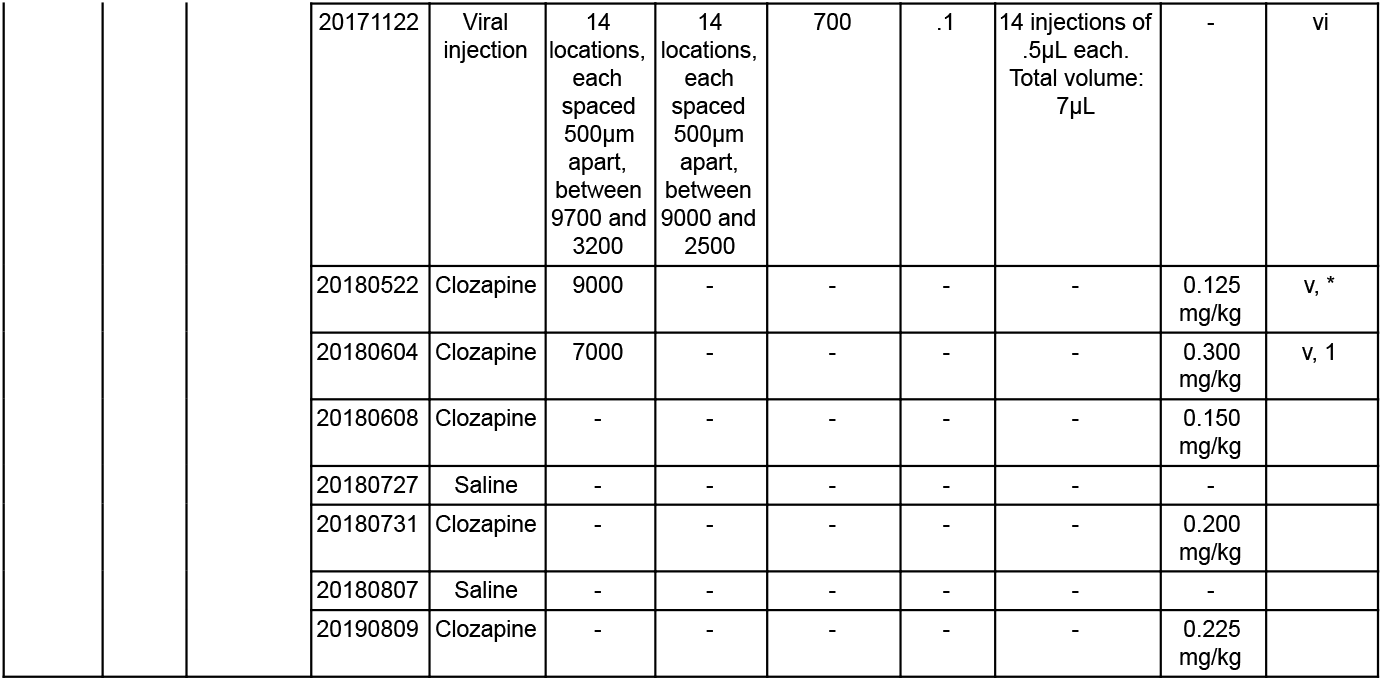
Session information List of all experimental sessions. Sessions are sorted by date for each monkey. The electrode and pipette depths are in micrometers below the dura. Electrode depth was constant throughout the session and we list the depth of either the tip of the electrode (single channel) or deepest electrode (24-channel V-probe). For muscimol infusion, the pipette was placed at four different depths. The injected volume is reported for each depth. For the muscimol infusion experiments, the distance between the recording electrode and the deepest pipette location is reported (i.e., first injection site). Symbols and abbreviations: Motion: Random-dot motion discrimination task. Time: Temporal-order discrimination task. *: Monkey completed < 500 trials post injection. %: Strongest motion strength (coherence +/-51.2%) was not used in this session. #: Low-volume muscimol injection session. !: Last session in Monkey 3. Further sessions were not possible due to the health and safety restrictions related to the COVID-19 pandemic. vi: Viral vector injection session. v: Sessions with V-probe recordings. Data shown in Figure 2D. N: Collected data on side-preference task. 1: First (high-dose) session. Data shown in Figure 3.

The location of the recording electrode varied across sessions but was always at a location on the lateral bank of the IPS with strong multi-unit neural activity (MUA) before inactivation. We quantified the MUA (Fig. 2C,D) as follows. The raw voltage signal (30 kHz sampling rate) was bandpassed between 300 and 6 kHz. The mean and standard deviation (*σ*) of the filtered signal in the time window 90 s before initiation of inactivation established a baseline for comparison. The raw MUA is defined as the frequency of positive crossings of threshold 3*σ* above baseline. Fig. 2C,D show the MUA normalized to the average MUA during the baseline epoch.

Injections were made with a Hamilton syringe (1700 series, gas tight, 50 μL volume) using a micro-injection pump (Phd Ultra-nanomite, Harvard Apparatus Inc.) connected to the pipette with Tygon tubing (0.25 mm inner diameter). The Hamilton syringe was filled with silicone fluid (Octamethyltrisiloxane; Clearco Products) mixed with fluorescent leak-detection dye (Dye-Lite; Tracerline) and filter-sterilized by passage through a Mixed Cellulose Esters membrane (Millex-GS 0.22 μm; SLGS033SB; EMD Millipore). The dyed silicone fluid allowed visualization of the meniscus to confirm the injected volume based on the length of travel of dye along the tubing.

We infused muscimol (8 μg/μL × 0.4 μL/min) at four depths along the injection track. The first of the four injections made during each session was at the deepest target location. After each injection, the pipette was left in place for at least 5 minutes before retraction to the next injection site. After each session, we confirmed that the pipette was intact by turning the pump on and visualizing a drop of fluid at the pipette tip. Table 3 shows injection details for the individual sessions. The total volume was typically 20 μL per session. However, in the first session (Monkey 1) the total volume was 45 μL, and in the fourth session the total volume was 8 μL. The low-volume injection failed to induce behavioral effects and we reverted to 20 μL in subsequent sessions. This session is excluded from the analysis in Figure 4A, but the reported effect is statistically significant with this data point included.

Saline injections followed the same injection protocol. We limited the number of saline injections to avoid tissue damage at the injection site (Zhou and Freedman, 2019). In sham sessions, all procedures were identical to those used in the muscimol and saline injections except that the pipette remained in the guide tube instead of being lowered into the brain, and the syringe was not connected to the pipette. In some sham sessions, we did not lower the electrode into the brain. We refer to both saline injections and sham sessions as control sessions.

### Chemogenetic inactivation and neural recordings

In Monkey 4, we injected the viral vector AAV5-hSyn-hM4Di-mCherry (titer = 4.9 × 10^12^ genome copies/mL, RRID: Addgene_50475) at locations informed by the mapping experiments (Fig. 2B). Injection procedures were similar to those described above for drug injections. The differences are detailed here. The viral vector was administered with a custom injectrode, comprising a pipette affixed to an electrode that protruded 700–800 μm beyond the tip of the pipette. The injectrode was lowered into the brain through a single transdural guide tube using a motorized hydraulic drive (FHC Inc.). Before injecting, we confirmed that the injectrode was at a location where neurons showed persistent activity during saccadic tasks. Injections were made along two tracks, separated by 1.4 mm, on two consecutive days. Each day, we injected at 13–14 depths separated by 500 μm covering 5.5–6mm. We injected 0.5 μL at each location at a rate of 0.1 μL/min, starting at the deepest location. The total injected volume was 13.5 μL. After each injection, the injectrode was left in place for an additional 8 minutes before being retracted to the next site. We then waited 6 months for expression of the hM4Di receptor to stabilize before beginning behavioral experiments.

In the inactivation experiments, we administered the hM4Di agonist, clozapine (Hello Bio #HB1607, concentrations listed in Table 3). Clozapine was chosen over the designer drug CNO as it is a more potent agonist of hM4Di receptors in the central nervous system at doses less than 10% of the minimum dose used clinically (Gomez et al., 2017; Raper et al., 2017). The monkey was trained to present its right arm through an opening in the primate chair to allow for subcutaneous clozapine injection. During two inactivation sessions, we recorded the effect of clozapine administration on neural firing rate with 24-channel V-probes (Plexon Inc.). The V-probe recordings were made 1–1.4 mm from the the viral injections. Following the session in which clozapine was administered at 0.15 mg/kg (see Table 3), Monkey 4 lost the cranial implant that allowed head stabilization. Subsequently, we were able to collect data from two additional inactivation sessions and two control sessions using a noninvasive restraint system.

### General procedures

During experimental sessions, the recording electrode was lowered into the brain and left in place until the end of the session. Baseline behavioral data were collected for at least 200 trials of the relevant task (motion direction or temporal order task). For Monkey 1 and 4, we then initiated the relevant inactivation procedure. For Monkey 2 and 3, we used inclusion criteria based on psychometric data to decide whether the behavior was sufficiently stable to continue the experiment. We computed the subjective point of equality (SPE) from logistic fit to the choice data (−*β*_1_/*β*_0_ from Eq. 2). Monkey 2 would continue the session only if |SPE| ≤ 3.2% coherence and the error rate at the highest coherence was ≤ 5%. For Monkey 3 the criteria were |SPE| ≤ 0.026 seconds and error rate ≤ 5%. Based on these inclusion criteria, we aborted 7 sessions for Monkey 2, and 7 sessions for Monkey 3.

During muscimol administration, the animals watched cartoon movies and received occasional juice rewards for looking at the screen. The pipette was left in place for at least 15 minutes afterwards, and behavioral data collection resumed after pipette removal. In the chemogenetic inactivation sessions, the animal waited for at least 30 minutes after clozapine administration before the collection of behavioral data resumed. On most sessions, monkeys performed at least 500 trials following inactivation. These 500 trials were included in the post-inactivation analysis. At the conclusion of each session and after removing the electrode and pipette from the chamber, we collected data in the extinction task (Monkeys 2 and 3 only). After each inactivation session, the animal did not work on any task for at least 3 days.

### Behavioral data analysis

We analyzed the effect of inactivation on choice using a variety of generalized linear models (GLM; logistic regression). The simplest generates the fits in Figure 3.

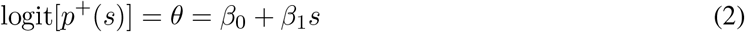

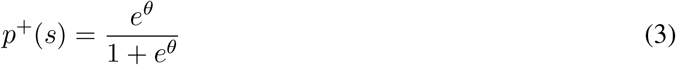

where *p*^+^ is the probability of a contralateral choice and *s* is the signed motion coherence (motion direction task) or the signed Δ*t* (temporal order task). In all cases, *s* > 0 indicates support for the choice target in the hemifield contralateral to the site of inactivation.

In the motion task, the strength of motion is a function of the coherence (coh) on the RDM stimulus and the duration of the presentation, *t*. The strength of the stimulus is therefore captured by a power law

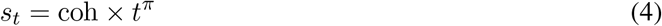

For perfect, unbounded accumulation of independent samples, the exponent would be *π* = 0.5, (i.e., the rate of improvement of signal to noise in the accumulation of independent identically distributed random samples), but the presence of a terminating bound attenuates the improvement (Kiani et al., 2008). The exponent used here was derived by fitting Eq. 2, with *s* = *s_t_*, to the control data (*π* = 0.38 and 0.43 for Monkeys 1 and 2, respectively). Using pre-injection data from all sessions, we confirmed that the version of Eq. 2 with *s_t_* is superior to a model that ignores stimulus duration (ΔBIC=31 for Monkey 1 and 27 for Monkey 2; *strong* support; Kass and Raftery 1995). We use Eq. 4 for all statistical analyses of the motion experiments. For the asynchrony experiment *s_t_* = Δ*t* = *s*, as defined above. Significance tests are standard t-tests, based on the standard error of the parameter, or *χ*^2^-tests, based on the difference in the deviance of nested models with and without the terms that define the null hypothesis, 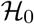.

For comparing inactivation-induced bias to pre-existing bias (in the same session) or to the bias on comparable trials during control sessions, we used the GLM,

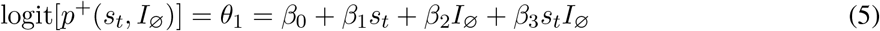

where *I*_∅_ = 1 if the trial occurs after administration of muscimol or clozapine, and 0 otherwise. To test whether inactivation produces a bias against contralateral choices, the null hypothesis is 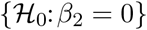. The curves shown in Fig. 3A use the expectation of *s_t_*:

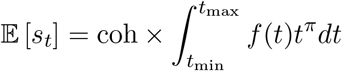

where *f*(*t*) is the distribution of durations defined in Eq. 1 and *π* takes the values defined above.

### Change in bias across sessions

To visualize compensation across sessions in Monkeys 1–3 (Fig. 4A) and across clozapine dosage in Monkey 4 (Fig. 4B) we used the GLM:

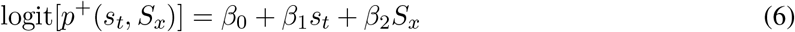

where *S_x_* is either the *x^th^* session number in chronological order (for Monkeys 1–3) or the dose of administered clozapine in mg/kg (for Monkey 4). For this analysis we use only the first 100 trials after inactivation. Lines in Fig. 4A–B are from this fit as are the *p*-values reported in Results. We confirmed that the effect of session number (or clozapine dose) on behavior is statistically significant even when the following saturated model was considered:

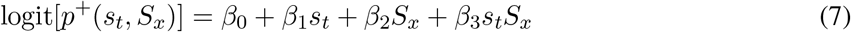

### Change in bias during a session

To visualize the decay of the choice bias over the course of a session (Fig. 4C–D), we compared *β*_0_ terms for Eq. 2, computed from trials 1–100 and from trials 401–500 (or the last 100 trials if the animal did not complete a 500 trial block after inactivation). Due to compensation across sessions, we could not detect a bias post-inactivation in some of the later sessions. We therefore only analyze sessions in which there was a significant bias in the first 100 trials. To compute the rate of compensation across trials in individual sessions, we added the term *N*_∅_, the trial number after inactivation, to Eq. 2:

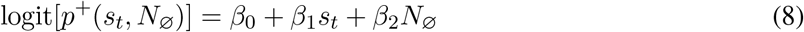

Finally, the statement about the effect of inactivation on sensitivity (Monkey 1) is supported by combining the first three inactivation sessions and elaborating Eq. 5 to include the trial number (post-injection) in each session:

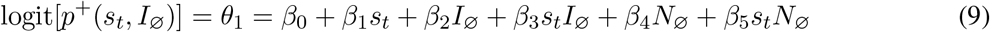

We report the p-value associated with 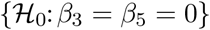, using the last 100 pre-injection trials and the first 500 post-injection trials from each session.

### Histology

We verified expression of the hM4Di-receptor in Monkey 4 histologically. The animal was euthanized under deep isofluorane anesthesia and perfused transcardially with 4% paraformaldehyde followed by a gradient of sucrose in phosphate buffer (10, 20, and 30%). The brain was extracted and cryoprotected in 30% sucrose. Sagittal sections (50 *μ*m) were cut on a sliding microtome and mounted onto slides. Transduced cells were first localized by inspecting native fluorescence signals. Sections were then stained using primary antibodies against the reporter proteins mCherry (Clontech 632543 RRID: AB_2307319, 1:250) and against the panneuronal marker NeuN (millipore MAB377 RRID: AB_2298772, 1:250), and using secondary antibodies (Invitrogen Molecular Probes): Alexa 568 (A10042 RRID: AB_2534017, 1:400), Alexa 488 (A21206 RRID: AB_141708 and custom, 1:400) and the nuclear stain DAPI (Invitrogen Molecular Probes D-21490, 1:5000) for visualization by epifluorescence microscopy.

### Data availability

Matlab code (m-files) and data (mat-files) to generate the main figures and for performing statistical analysis will be made available in a public github repository at the time of publication.

---

**Supplementary Figure 1.**
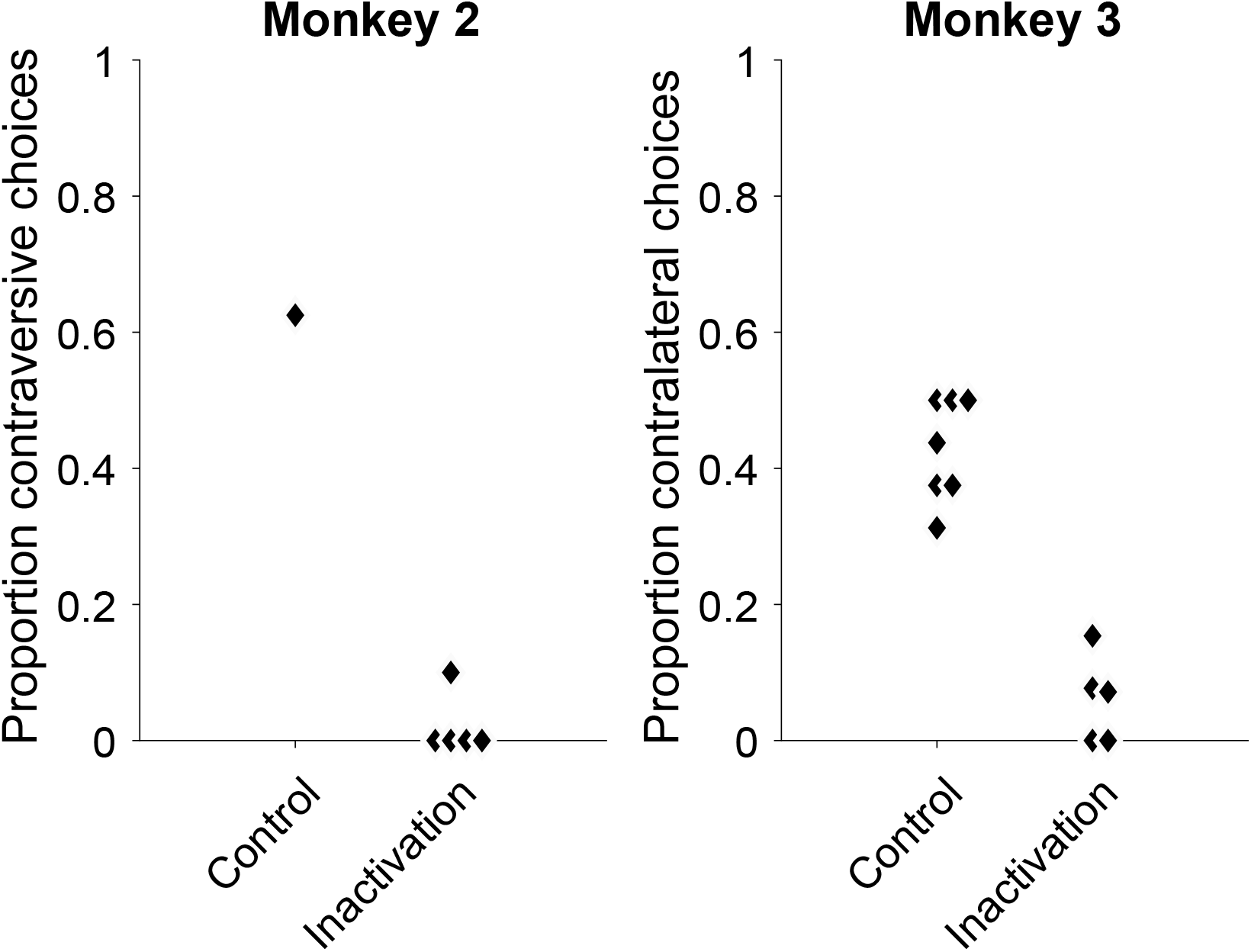
Behavior on the side-preference task. After completion of the main experiment, Monkeys 2 and 3 were tested on a preference task intended to approximate a neurological double-simultaneous stimulation test for extinction. The experimenter presented desirable food items in both palms symmetric about either side of the monkeys mouth, and allowed the monkey to choose and item by picking the treat with its tongue. The proportion of chosen items from the side contralateral to the inactivated cortex is shown. Points are data from one session. Points belonging to the same treatment group (control or muscimol inactivation) are displaced horizontally for visualization.

**Supplementary Figure 2.**
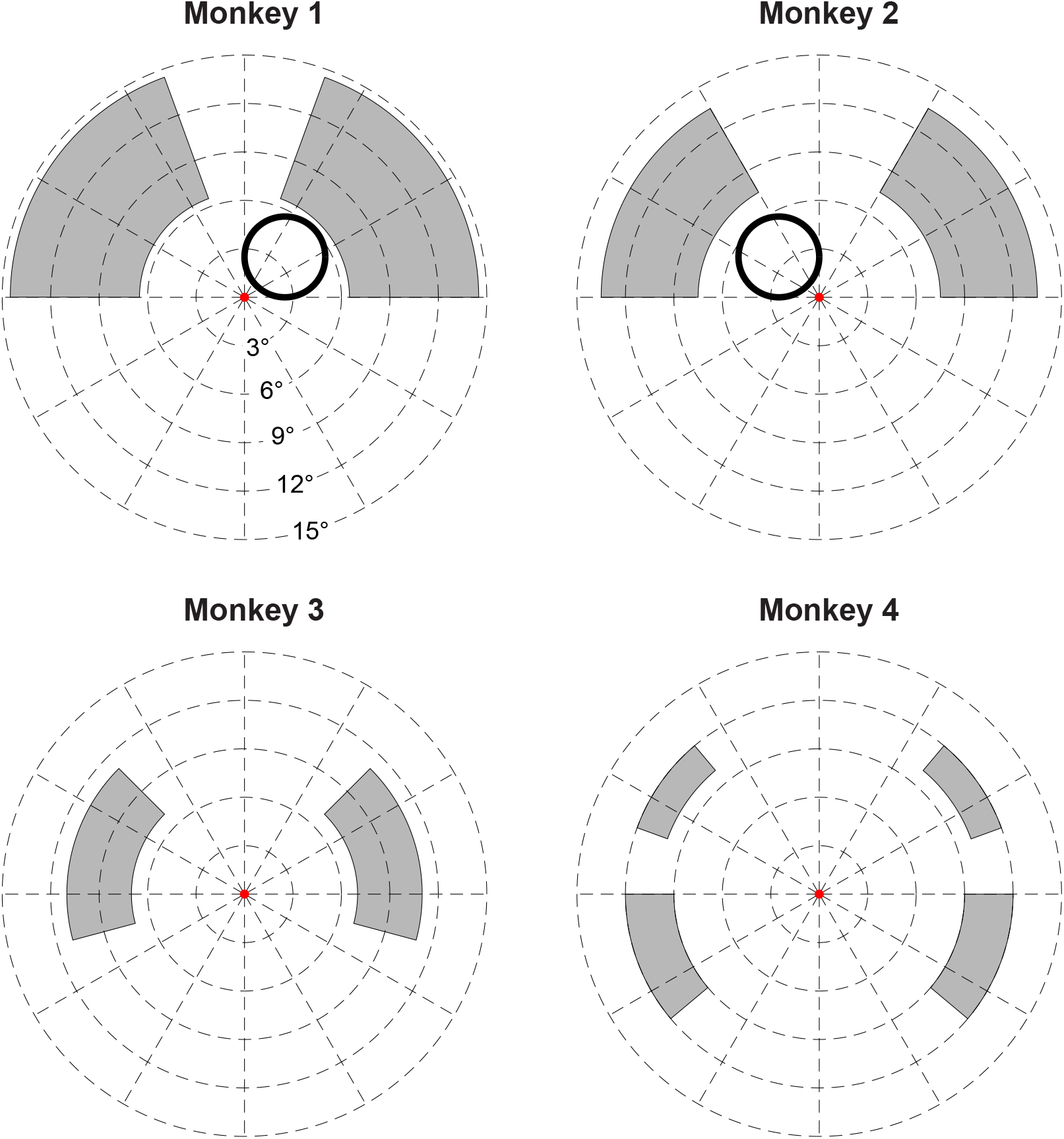
Stimulus configuration. The left and right choice-targets were positioned randomly on each trial using independent samples from the shaded regions of the visual field (uniform distribution over range of *r*,*θ*). For monkey 4, both choice targets were in either the upper or the lower hemifield. The area subtended by the random dot motion display (black circles; Monkeys 1 and 2) was consistent across trials/sessions and confined to the hemifield ipsilateral to the inactivated cortex. Eccentricities are in degrees visual angle.

## Notes

**Conflict of interest statement:** The authors declare no competing financial interests.

### Competing Interest Statement

The authors have declared no competing interest.

